# OCT5k: A dataset of multi-disease and multi-graded annotations for retinal layers

**DOI:** 10.1101/2023.03.29.534704

**Authors:** Mustafa Arikan, James Willoughby, Sevim Ongun, Ferenc Sallo, Andrea Montesel, Hend Ahmed, Ahmed Hagag, Marius Book, Henrik Faatz, Maria Vittoria Cicinelli, Amani A Fawzi, Dominika Podkowinski, Marketa Cilkova, Deanna de Almeida, Moussa Zouache, Ganesham Ramsamy, Watjana Lilaonitkul, Adam M Dubis

## Abstract

The thickness and appearance of retinal layers are essential markers for diagnosing and studying eye diseases. Despite the increasing availability of imaging devices to scan and store large amounts of data, analyzing retinal images and generating trial endpoints has remained a manual, error-prone, and time-consuming task. In particular, the lack of large amounts of high-quality labels for different diseases hinders the development of automated algorithms. Therefore, we have compiled 5016 pixel-wise manual labels for 1672 optical coherence tomography (OCT) scans featuring two different diseases as well as healthy subjects to help democratize the process of developing novel automatic techniques. We also collected 4698 bounding box annotations for a subset of 566 scans across 9 classes of disease biomarker. Due to variations in retinal morphology, intensity range, and changes in contrast and brightness, designing segmentation and detection methods that can generalize to different disease types is challenging. While machine learning-based methods can overcome these challenges, high-quality expert annotations are necessary for training. Publicly available annotated image datasets typically contain few images and/or only cover a single type of disease, and most are only annotated by a single grader. To address this gap, we present a comprehensive multi-grader and multi-disease dataset for training machine learning-based algorithms. The proposed dataset covers three subsets of scans (Age-related Macular Degeneration, Diabetic Macular Edema, and healthy) and annotations for two types of tasks (semantic segmentation and object detection).

## Background & Summary

Ophthalmologists use retinal images to track disease progression, analyze disease patterns, approach diagnosis and monitor treatment^1–3^. The availability and mass usage of OCT devices made them the gold standard for imaging in ophthalmology^4^. Retinal imaging using OCT has seen major advancements in image resolution and acquisition speed. Experts are now in need of fast and robust models for analyzing large amounts of images and making sense of them. For example, AMD (Age-related Macular Degeneration) is a disease that affects the macular area of the retina and it is the leading cause of irreversible blindness and severely impaired eyesight. There were 170 million patients with AMD in 2020 and this number is increasing^5^, this is resulting in larger quantities of imaging data needing to be analyzed by ophthalmologists to keep up with the increase in patients.

The measurement of the retinal layers is a critical step in the analysis of eye diseases^6–10^ and as such automated systems to assist with this measurement are very desirable. Images of the retinal layers are obtained by OCT devices. Retinal layers can be annotated manually, semi-automatically, or automatically and the thicknesses can then be calculated using segmentations. As a result, the changes in thickness between retinal layers can be used for analyzing and modeling the progression of diseases^11^. In a manual setting, the layers of the retina are delineated using lines to distinguish distinct areas of the retina. OCT volumes that need to be annotated can contain more than 100 images per patient and per scan, which increases the amount of work and makes manual annotation a tedious and time-consuming process.

For the quantitative analysis of large amounts of scans to be reliable, accurate and robust segmentation of retinal layers in OCT images is necessary. For handling large amounts of imaging data and making statistical analysis sufficiently powerful, fully automated segmentation algorithms^12–16^ are preferred. Although images representing retinal layers for a single disease can be segmented automatically with fewer images and annotations for training, more sophisticated algorithms are required to segment retinal layers for a wider set of diseases for better generalization. As a result, publicly available databases and platforms which offer researchers access to image datasets and annotations that can be used to produce sophisticated algorithms for complex automatic ophthalmic image analysis are very desirable^17^.

In an automatic setting, obtaining retinal thickness measurements can be divided into two separate tasks. The first step involves assigning labels to each pixel in the image. This procedure is referred to as semantic segmentation. In the second step, the segmentation outputs are used to identify the lines that delineate the layers within the retina. Therefore, the obtained lines represent the borders of the retinal layers, which can then be used to calculate and store thickness measurements for further analysis.

Publicly available data sets in combination with artificial intelligence methods like deep learning have great potential to transform research on eye diseases and retinal imaging^18^. A major challenge in deep learning is the need for high-quality labels^19^. To train accurate models and compare different algorithms against one another and against expert grading, the quality of the labels is crucial. Therefore, in order to understand the uncertainty^20^ properties of data and models, we need to collect annotations for the same images from different graders.

Publicly available OCT datasets for retinal layer segmentation have been limited in scope, often being small in size, specific to a single disease, or containing only one grading. It is worth mentioning that one major limitation of such datasets is the lack of power to detect less common phenotypes, which may impact the performance of segmentation algorithms on rare diseases and diverse patient populations. Li et al.^21^ introduced a data set for the glaucoma disease for which there was a single annotation for 9 layers and 122 peripapillary OCT scans. The data set was augmented with flips to increase its size to 244 images. Hassan et al.^22, 23^ published a dataset with 42 scans and 6 layers annotated. In He et al.^24^, 1715 annotations, composed of 8 layers each, were collected for 14 healthy and 21 multiple sclerosis OCT volumes. As part of a study published in Anthony et al.^25^, layer annotations were manually collected from three graders on some 40 mouse OCT slices. The data set also includes automatic segmentation of 6 and 10 layers for 40 dense OCT volumes. In Gholami et al.^26^ manual annotations for 25 healthy scans are published. Farsiu et al.^27^ published 269 AMD volumes and 115 control volumes with three layers, which were semiautomatically segmented. In Tian et al.^28^, the authors have compiled a dataset containing annotations for 100 individual OCT slices taken from healthy individuals. Chiu et al.^29^ collected two sets of manual gradings for 110 OCT slices and 8 retinal layers. In Chiu et al.^30^, they collected two sets of manual gradings for 220 OCT slices and three retinal layers. Melinscak et al.^31^ published a dataset for AMD containing 1136 images annotated with 3 types of fluids and 4 retinal layers. An additional annotation was made for a subset of 75 OCT slices in order to calculate intra-observer and inter-observer error. Morales et al.^32^ have collected data for 6 retinal layers and 244 slices from rat OCT volume data.

To address the limitations of existing data sets, we have provided a large set of annotations for 60 OCT volumes - 20 from individuals with AMD, 20 from individuals with diabetic macular edema (DME), and 20 from healthy individuals – taken from a pool of 148 OCT volumes^33^. The volumes were obtained using a Heidelberg Spectralis (Heidelberg Engineering, Germany) device. The axial resolution is 3:5.m and the scan-dimension is 8:9.7:4 mm2. The number of scans per volume varies between 19 and 69 scans. A complete list of volumes per disease/group is given in Table 1.

**Table 1.** The total number of individual scans that were graded manually by three different annotators is 1672.

| Disease/Group | Number of total scans | Number of volumes x scans per volume |
| --- | --- | --- |
| AMD | 722 | 7 x 19, 4 x 25, 2 x 31, 7 x 61 |
| DME | 422 | 19 x 19, 1 x 61 |
| Healthy | 530 | 12 x 19, 2 x 25, 3 x 31, 1 x 37, 2 x 61 |

Additionally, 4698 bounding box annotations for a subset of 566 scans are provided across nine classes. These 9 classes are choroidal folds, retinal fluid, geographic atrophy, hard drusen, hyperfluorescent spots, photoreceptor layer disruption, reticular drusen, soft drusen, and softdrusen PED (Pigment Epithelial Detachment). We have also included automatic retinal layer segmentations for the remaining 88 OCT volumes, giving a total of 2586 annotations.

In conclusion, we propose a set of multi-graded, multi-disease annotations for a heterogeneous real-world dataset that consists of pixel-wise annotations for retinal layers, as well as bounding box annotations for different classes of biomarkers in OCT images. It can be used to train and evaluate machine learning-based retinal layer segmentation architectures, and to test their ability to generalize to different disease settings.

## Methods

We first describe the annotation process of the retinal layers. In the subsection, titled “Inter-grader Reliability”, we present the details of the process preparing multiple gradings for the same scans. In the second subsection, titled “Object Detection Labels”, we present the details of the bounding box annotations. In the section titled “Segmentation Model”, we describe the baseline method used to segment the retinal layers.

### Ground Truth Annotation

The ground truth annotation involves human-machine interaction, tooling, finding best practices for annotation and infrastructure and data management. We have built a customized labeling platform based on the Hitachi’s open source semantic segmentation editor^34^ to allow multiple graders to annotate OCT images. The OCT images are uploaded to the platform and the layers are drawn according a defined layout. The layout defines the number of layers and the colors assigned to the layers. The layout is adjustable, therefore the number of layers can be changed. The number of layers depends on disease and corresponding thickness of certain layers.

The images for annotation were prepared, shuffled and uploaded to the labeling platform. The graders received access to a portion of the images. To speed up annotation we trained a model using a small number of images. We used this model to prepare segmented masks of retinal layers, which were uploaded as well. The annotation masks were replaced by lines and vertices which delineated the layers. The vertices can be dragged and dropped to change the annotation. In addition the vertices can be deleted and new ones can be added. The graders navigated through folders and images and corrected the five retinal layers. To reduce bias, the images from the same volumes were shuffled and distributed among graders. After the correction of the retinal layers the results were compiled. In a second stage outliers were determined and checked and if necessary corrected again. For the outlier detection we used the interquartile range (IQR). We repeated this step three times and and checked the quality of the layer annotations and improved it iteratively.

The agreement or disagreement between different expert graders is valuable for model development and for highlighting uncertainty and providing feedback to experts. To ease this task and to support further development of uncertainty in model and data we compiled three sets of annotations with help from different graders.

All graders performed the annotations according following guidelines. A sample workflow is visualized in 1.

- ILM: This layer is the furthest from the top and should be segmented by following the pattern of the band.
- OPL: OPL is below ILM and is segmented on top of the bright band of the OPL.
- IS/OS: IS/OS layer could be segmented by distinguishing that the IS/OS boundary would be located above the RPE. This layer may also be referred to as EZ (ellipsoid zone).
- OBRPE: Segmenting the RPE (top and bottom): The outermost highly reflective layer is called the Outer boundary of RPE which is segmented along the bottom band of the layer.
- IBRPE: Inner boundary of RPE is segmented along the top band of the Outer Boundary of RPE layer.

The layout of the annotation platform can be easily adjusted and expanded for different retinal layers. This is necessary, since different retinal layers are effected by different eye diseases. Thus, we can use the annotation platform to annotate different OCT scans from different diseases. In figure 1 a layout for manual annotation is presented.

**Figure 1.**
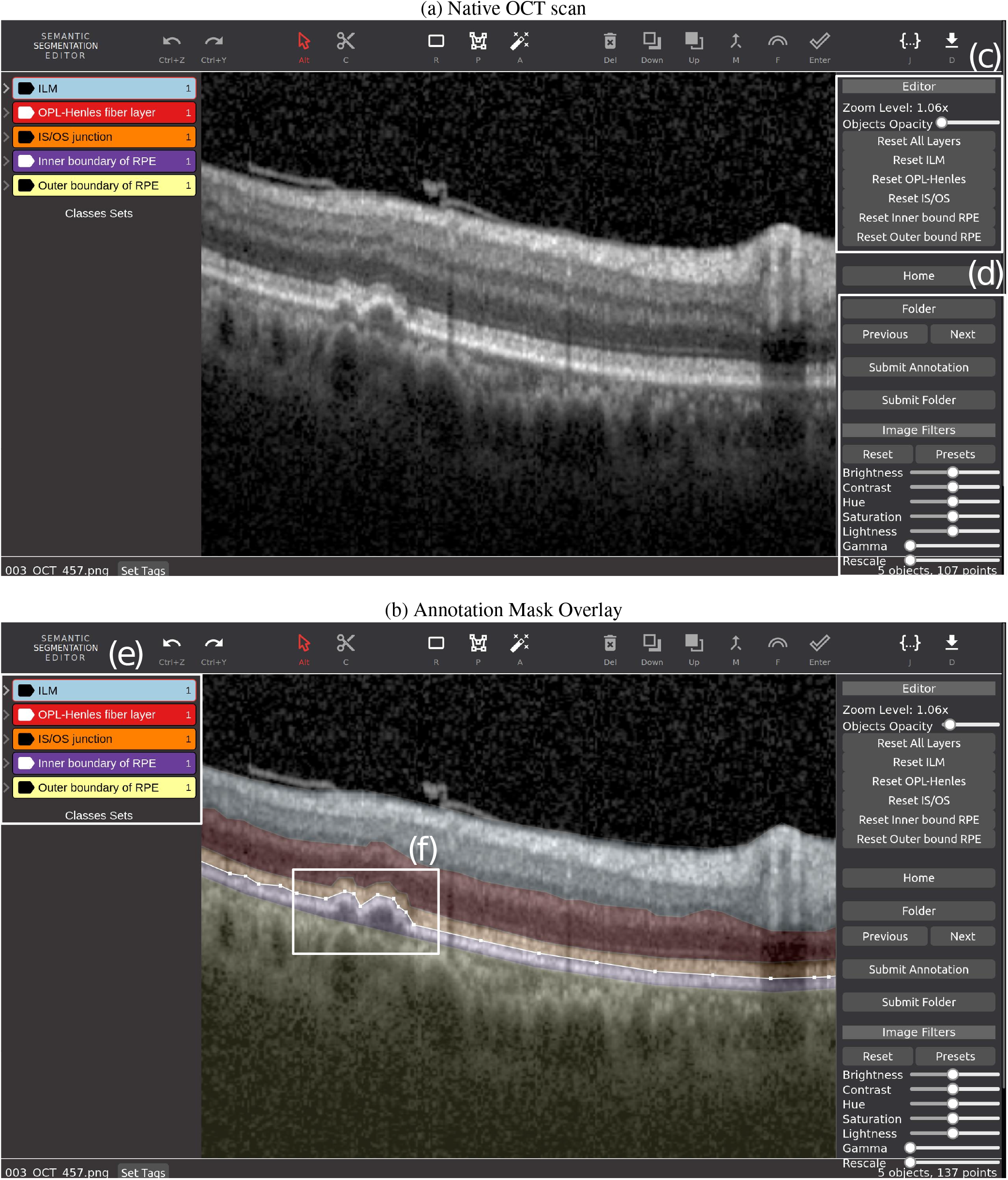
(a) The annotation platform with a layout to annotate five retinal layers, (b) and with opacity adjusted, (c) functions to change opacity and resetting layers and (d) and buttons for navigating, submitting gradings and changing image visualization options such as contrast or hue. (e) Custom layout for five retinal layers, (f) Area showing editing using vertices.

For the manual annotation and correction of the data set we selected five retinal layers. The graders annotated Internal limiting membrane (ILM), Outer Plexiform Layer (OPL), Photoreceptor Inner Segment and Outer Segment Layers (IS-OS), Inner Boundary of Retinal Pigment Epithelium (IBRPE) and Outer Boundary of Retinal Pigment Epithelium (OBRPE) layers. The retinal layers with a corresponding image and annotation mask is presented in Figure 2 for AMD and DME.

**Figure 2.**
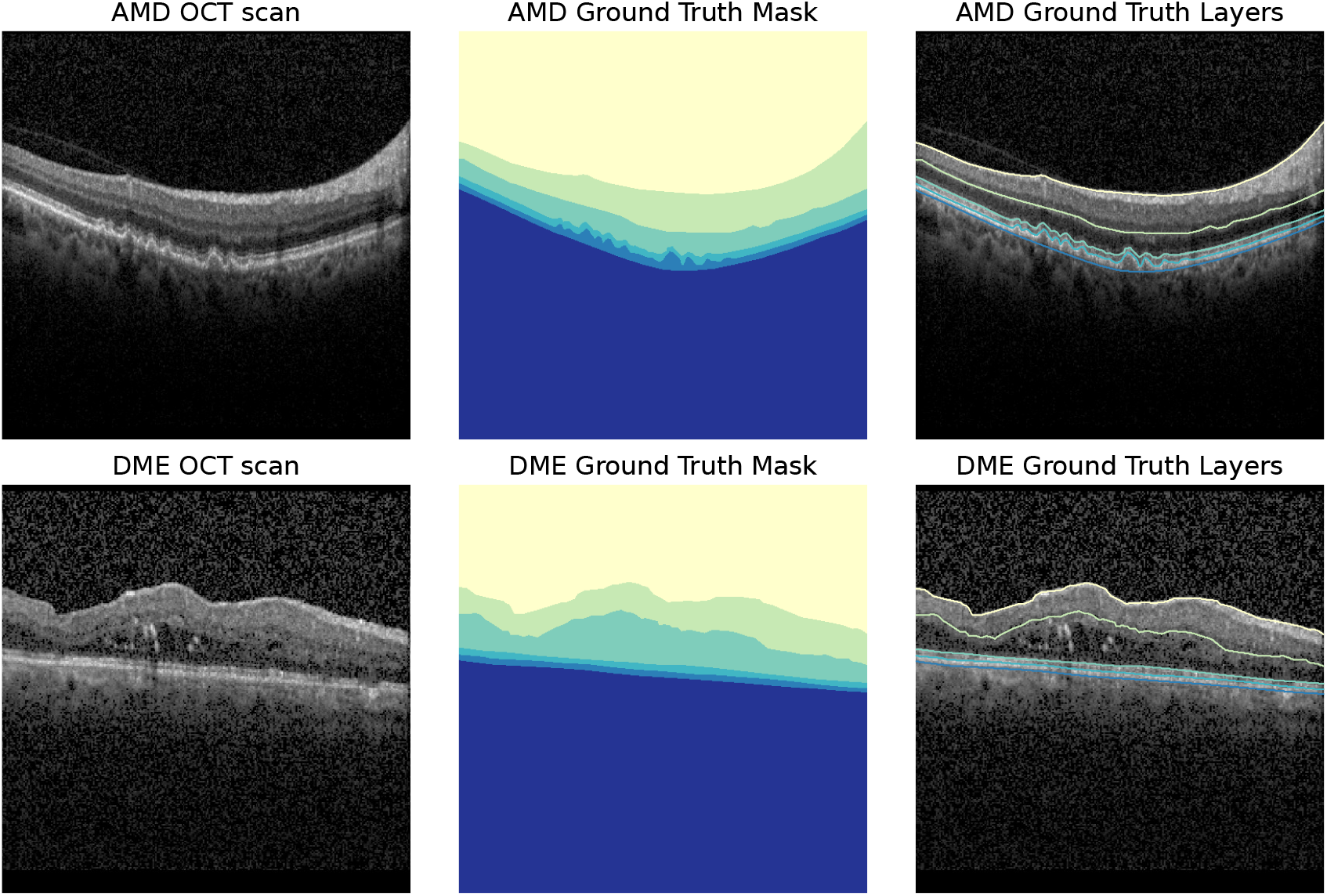
A single scan from AMD (top) and DME (bottom) (left) and the exported retinal layers from the labeling platform (right). The annotated retinal layers are (from top to bottom): ILM, OPL, IS-OS, IBRPE and OBRPE. And the generated masks for a semantic segmentation task (middle).

#### Inter-Grader Reliability

In order to evaluate the uncertainty in medical image segmentation, models based on uncertainty estimation with dropout or training ensembles have been examined. Furthermore, the determination of when results are to be considered uncertain is task- and data-dependent as well as the result of multiple grades from multiple graders. To date, there are approaches for creating large datasets with single annotations, but no large datasets exist for evaluating the accuracy of realistic uncertainties based on consensus reached from multiple gradings to determine which procedures for uncertainty estimation produce realistic results.

For the purpose of calculating inter-grader error, annotations for 1672 OCT scans were performed by three different sets of graders. Specifically, models for uncertainty estimation can be compared against uncertainties that human graders attribute to different retinal layers and structures. Thus, beside providing a large dataset for training novel layer segmentation models, we enable the development and evaluation of uncertainty estimation methods.

#### Object Detection Labels

Besides retinal layers, we have collected bounding box annotations^35^ for 9 different classes and a subset of 566 images. The bounding box annotations were created by a set of experts who were supplied with images from the AMD scans of the Rasti et al.^33^ and Kermany et al.^36^ datasets. Each expert was given a unique set of images to annotate with the exception of one set of 100 images which was annotated twice by separate experts. Geographic Atrophy bounding boxes were required to span the entire width of the transillumination defect with the inclusion of up to 5 pixels either side and were required to include the first choroidal vessels on the bottom and minimal pixels at the top. Drusen annotations were required to encompass the peak of the drusen, the Bruch’s boundary and both points where the drusen joins the RPE. Each drusen was annotated individually according to these requirements.

The bounding box annotations is composed of 566 Images with a total of 4698 objects distributed across nine classes. The distribution of the objects between the classes is given in the table 2 below. In figure 3 we present two examples of OCT scans containing bounding box annotations.

**Table 2.** The total number of individual bounding box annotations.

| Class | Number of Annotations |
| --- | --- |
| Soft Drusen | 1798 |
| Hard Drusen | 1204 |
| Photoreceptor layer disruption | 711 |
| Reticular drusen | 326 |
| Soft drusen&PED | 302 |
| Hyperfluorescent spots | 200 |
| Choroidal fold | 151 |
| Geographic atrophy | 128 |
| Fluid | 123 |

**Figure 3.**
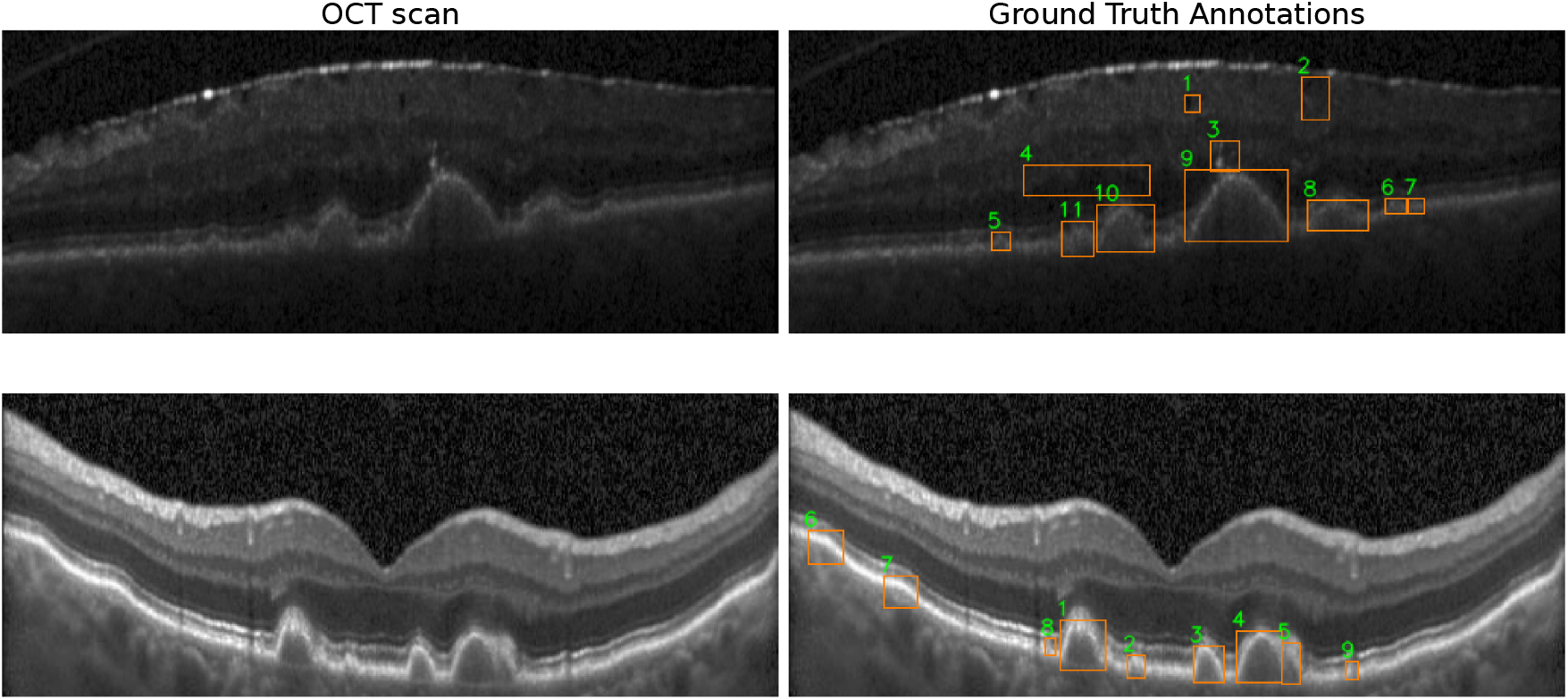
Examples from two scans with bounding box annotations: Top row has fluid (1,2,4), Hyperfluorescent spots (3), Harddrusen (5,6,7) and Softdrusen(8,9,10,11). Bottom row has Softdrusen PED (1,3,4,5), Softdrusen (2,8,9) and Choroidalfolds (6,7).

### Segmentation Model

Deep learning networks are used to solve certain problems depending on the scale of data (such as the amount, resolution, or number of channels). The networks are manually customized to achieve better accuracy for different tasks with larger data sets and/or images with higher resolution (for instance, by adding more layers).

For the semantic segmentation we use a U-Net with an EfficientNet backbone. EfficientNet^37^ proposes scaling up deep learning models in order to improve accuracy and efficiency much more easily. It uses a technique called compound coefficient scaling to modify each dimension uniformly with a fixed set of coefficients. Using this scaling method and AutoML (Automated machine learning), seven models were created, which outperformed most convolutional neural networks and achieved state-of-the-art performance. Thus, we choose EfficientNet to have the ability for training with different image resolutions and easier adoption to different modalities. Thus, we choose EfficientNet^38^ to have the ability for training with different image resolutions and easier adoption to different modalities.

## Data Records

The layer annotations are saved in three formats. First, in text files where every line corresponds to a single column of the scan. Second, in grey value image files where every pixel is assigned to a layer. Third, in RGB images, where every pixel is assigned an RGB triplet representing a layer. In addition to annotations we provide scripts to download the original images from Rabbani et al.^33^ and scripts to prepare the images and annotations for model training. The data set can be downloaded from (https://doi.org/10.5522/04/22128671.v1) for research purposes.

## Technical Validation

The annotation strategy for the segmentation data relies on four metrics for annotation quality. Different metrics highlight different aspects of the annotations. They have both advantages and shortcomings. We have used four metrics to analyze the differences between graders for quality assessment and control. Thus, we can apply outlier detection using any of the four metrics for quality assessment and for sending back images for correction.

- Dice similarity coefficient: this is a metric to measure the similarity of two segmentation masks. It can be written as,

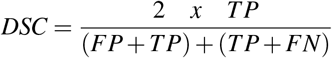

where TP denotes true positives, FP false positives and FN false negatives.
- Intersection over Union: the IoU metric is related to the DSC and it is calculated by the common pixels between the target and the prediction divided by the total number pixels present in both annotations or masks. The formula is:

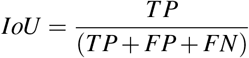
- Mean distance between lines: the mean distance between two lines representing layers is calculated by averaging the sum of absolute differences between two lines. The formula is:

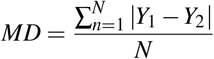

where N stands for number of pixels in x-direction and *Y*_1_ and *Y*_2_ stand for position of the layer in y-direction.

Intersection over Union (IoU) is a more robust metric for evaluating the semantic segmentation of retinal layers in OCT images compared to the Dice score. The main reason behind this is that IoU is less sensitive to class imbalance, which is critical for retinal layer segmentation due to the disparity in size between various layers. By calculating the ratio of the area of intersection between the predicted segmentation and the ground truth segmentation to their combined area (union), IoU provides a more accurate representation of the segmentation quality. IoU converges to zero when the ground truth and prediction have no overlap, while the Dice score may still provide non-zero values in such cases, further emphasizing the robustness of IoU for layer segmentation purposes. Therefore, IoU should be selected as the preferred evaluation metric for semantic segmentation of retinal layers in OCT images, since it ensures a more reliable and accurate assessment of the model’s performance.

Incorporating mean distance alongside IoU for evaluating semantic segmentation of retinal layers in OCT images addresses IoU’s shortcomings when layers disappear or become thin. Mean distance measures average boundary distances between predicted and ground truth layers, providing insight into localization accuracy. This additional metric helps detect systematic errors and offers valuable information on boundary consistency in extracted segmentation masks. By combining IoU and mean distance, a more robust and detailed evaluation of segmentation models handling complex tasks is achieved.

### Inter-grader Agreement

We used quality metrics to calculate the inter-grader agreement of annotations between multiple graders. Outliers in the annotations were detected and corrected in a second stage. We utilized the interquartile range (IQR) for outlier detection and iteratively improved annotation quality. Each grader had access to multiple images, and once they finished annotating, we compared their results using IQR to a second grader. Outliers were identified between two graders’ agreement on any of the five layers if they exceeded the IQR criteria. These outliers were then sent back to the graders for correction. This process was repeated three times to ensure high-quality annotations.

We evaluated the inter-grader agreement among annotations by examining both the Intersection over Union (IoU) and the mean distance metric across all six retinal bands. Using multiple metrics provides a more comprehensive quality assessment, as relying solely on the IoU for retinal layer segmentation might be insufficient due to potential misinterpretations that may arise from disappearing or thinning layers. In cases where two layers become connected as a result of disappearing or thinning layers, the IoU metric may yield a low value, indicating low agreement. However, such low agreement can be attributed to the absence of pixels in the band for comparison, rather than genuine disagreement among the graders. Thus, it is essential to employ multiple metrics to efficiently evaluate the quality of the dataset.

The quality of annotations between each pair, for the entire dataset as well as for specific groups (AMD, DME, and healthy), is illustrated in Figure 4. We visualized the IoU scores for inter-grader agreement among gradings 1, 2, and 3 across all six retinal bands. These bands include: Background/ILM, ILM/OPL, OPL/IS-OS, IS-OS/IBRPE, IBRPE/OBRPE, and OBRPE/Background. By combining these metrics, we have established a reliable and accurate evaluation of the annotations, suitable for academic publication.

**Figure 4.**
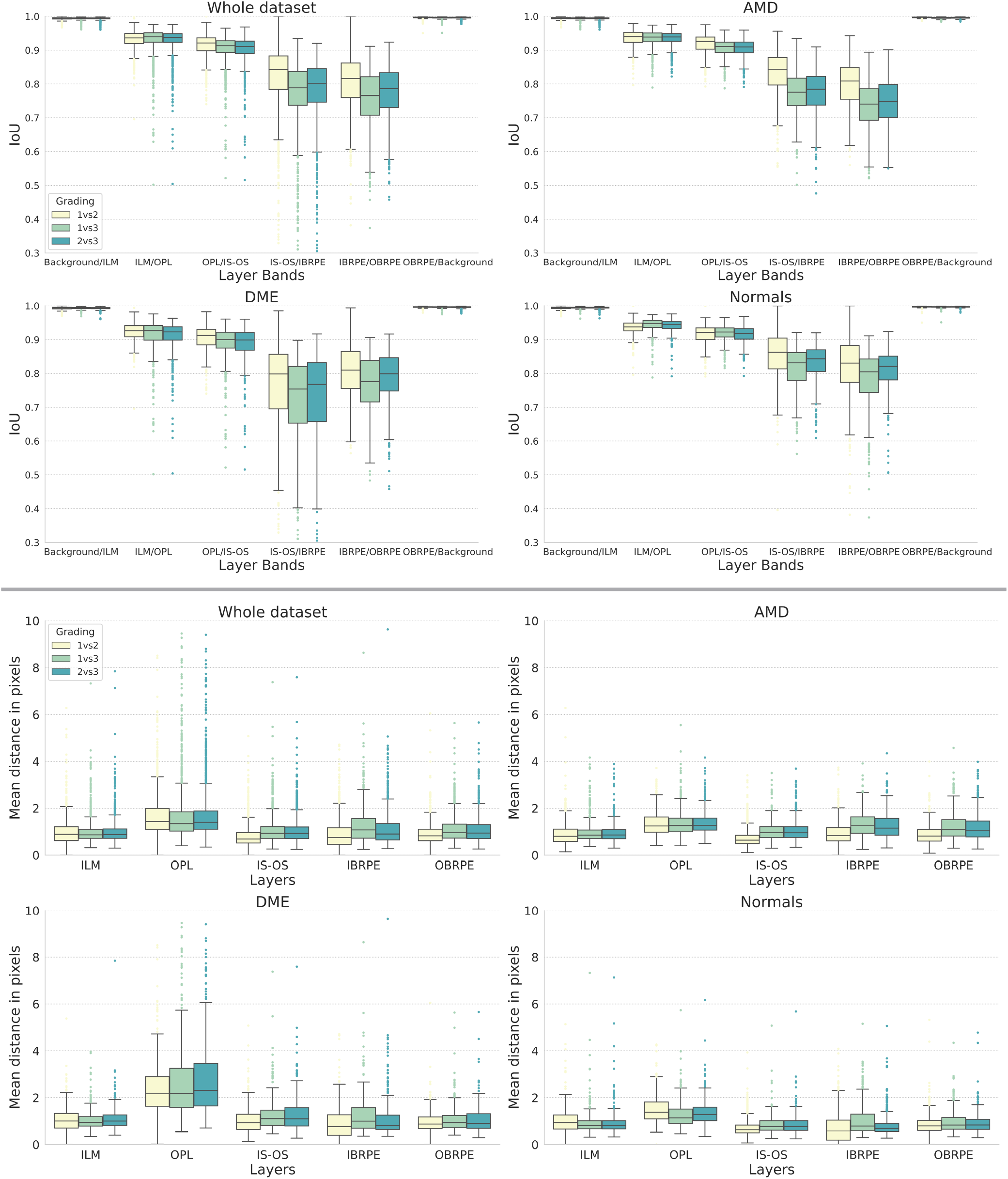
This figure shows boxplots for two comparison metrics, Intersection over Union (IoU) and Mean Distance, across the four categories of retinal images: AMD, DME, healthy, and the entire dataset. The top row displays IoU results for the whole dataset (top left), AMD (top right), DME (bottom left), and healthy (bottom right). The bottom row displays Mean Distance results for the same categories. Each boxplot compares two gradings for a particular annotation band, providing insight into inter-rater agreement for the dataset.

In the annotation process, comparing mean distances between lines representing layers offers an additional metric as it accounts for disappearing layers or those connected to either the top or bottom of the neighbouring layer. This requires the use of a post-processing method to maintain the continuity of disappearing layers, ensuring that they appear on the following retinal layer. By combining these metrics and our iterative inter-grader agreement process, we achieved high-quality annotations suitable for training and evaluation of our segmentation method.

## Usage Notes

We provide ground truth annotations in three different types. First, as layer annotations with pairs of x and y values representing the layers in pixel coordinates. Second, as annotation masks, where every pixel is assigned a grey value representing a band. The values are between 0 and 5. Third, as annotations masks with RGB values assigned to every pixel. The first type can be used for training sequence based models. The second type can be used for training semantic segmentation models (e.g. U-Net based). The third type is similar to the second type but it is intended for visualization and analysis purposes. The data set consists of annotations only, the original images are downloaded from external data sets. We provide scripts to prepare and combine both annotations and images.

## Code availability

Source code for data preparation is available within the data set file. It contains the following scripts:

- Prepare_Images_Download.ipynb: Code for downloading original images and preparing the folder structure.
- Prepare_Data_For_Model_Development.ipynb: Code for generating NumPy arrays containing annotations and images for model training.
- Prepare_Data_For_Detection.ipynb: Code for reading and visualizing bounding box annotations.

Detailed descriptions are in the README.txt within the data set file. A Python environment is needed to be able to run the scripts. We also provide a description for creating a virtual environment. The file “requirements.txt” contains all relevant software libraries and packages with corresponding versions.

## Acknowledgements

The authors acknowledge the use of the Joint Academic Data science Endeavour (JADE) Tier 2 computing facility funded by the Engineering and Physical Sciences Research Council (EPSRC), UK. The research leading to these results has received support from the UCL Institute of Healthcare Engineering (IHE) and the UCL Institute of Ophthalmology (IoO).

## Author contributions statement

MA built the annotation platform, compiled the data set and developed the baseline segmentation method. JW prepared the object detection labels and reviewed the manuscript. WL conceived the annotation procedure to reduce bias, helped developing the baseline method and reviewed the manuscript. AD planned the overall annotation, monitored the annotation process, selected OCT volumes and reviewed the manuscript. SO, FS, AM, HA, AH, MB, HF, MC, AF, DP, MC, DA, MZ and GR annotated scans, gave feedback for improving the platform and the manuscript.

## References

1. Imaging of macular diseases with optical coherence tomography. Ophthalmology 102, 217–229, https://doi.org/10.1016/S0161-6420(95)31032-9 (1995).

2. Jaffe, G. & Caprioli, J. Optical coherence tomography to detect and manage retinal disease and glaucoma. Am. journal ophthalmology 137, 156–69, 10.1016/S0002-9394(03)00792-X (2004).

3. Adhi, M. & Duker, J. Optical coherence tomography – current and future applications. Curr. opinion ophthalmology 24, 213–21, 10.1097/ICU.0b013e32835f8bf8 (2013).

4. Drexler, W. & Fujimoto, J. G. State-of-the-art retinal optical coherence tomography. Prog. Retin. Eye Res. 27, 45–88, https://doi.org/10.1016/j.preteyeres.2007.07.005 (2008).

5. Xu, X. et al. Regional differences in the global burden of age-related macular degeneration. BMC Public Heal. 20, 10.1186/s12889-020-8445-y (2020).

6. van Dijk, H. et al. Selective loss of inner retinal layer thickness in type 1 diabetic patients with minimal diabetic retinopathy. Investig. ophthalmology visual science 50, 3404–9, 10.1167/iovs.08-3143 (2009).

7. Acton, J., Smith, R., Hood, D. & Greenstein, V. The relationship between retinal layer thickness and the visual field in early age-related macular degeneration. Investig. ophthalmology visual science 53, 10.1167/iovs.12-10361 (2012).

8. Budenz, D. L. et al. Determinants of normal retinal nerve fiber layer thickness measured by stratus oct. Ophthalmology 114, 1046–1052, https://doi.org/10.1016/j.ophtha.2006.08.046 (2007).

9. Yanni, S. et al. Normative reference ranges for the retinal nerve fiber layer, macula, and retinal layer thicknesses in children. Am. journal ophthalmology 155, 10.1016/j.ajo.2012.08.010 (2012).

10. Fisher, J. B. et al. Relation of visual function to retinal nerve fiber layer thickness in multiple sclerosis. Ophthalmology 113, 324–332, https://doi.org/10.1016/j.ophtha.2005.10.040 (2006).

11. Podkowinski, D. et al. Neuroretinal atrophy following resolution of macular oedema in retinal vein occlusion. Br. J. Ophthalmol. 103, 36–42, 10.1136/bjophthalmol-2017-311614 (2019). https://bjo.bmj.com/content/103/1/36.full.pdf.

12. Gu, Z. et al. Ce-net: Context encoder network for 2d medical image segmentation. IEEE Transactions on Med. Imaging 38, 2281–2292, 10.1109/TMI.2019.2903562 (2019).

13. Fang, L. et al. Automatic segmentation of nine retinal layer boundaries in oct images of non-exudative amd patients using deep learning and graph search. Biomed. Opt. Express 8, 2732, 10.1364/BOE.8.002732 (2017).

14. Guha Roy, A. et al. Relaynet: Retinal layer and fluid segmentation of macular optical coherence tomography using fully convolutional network. Biomed. Opt. Express 8, 10.1364/BOE.8.003627 (2017).

15. Chiu, S. et al. Automatic segmentation of seven retinal layers in sdoct images congruent with expert manual segmentation. Opt. express 18, 19413–28, 10.1364/OE.18.019413 (2010).

16. Pekala, M. et al. Deep learning based retinal oct segmentation. Comput. Biol. Medicine 114, 103445, 10.1016/j.compbiomed.2019.103445 (2019).

17. Tong, Y., Lu, W., Yu, Y. H. & Shen, Y. Application of machine learning in ophthalmic imaging modalities. Eye Vis. 7 (2020).

18. Schmidt-Erfurth, U., Sadeghipour, A., Gerendas, B. S., Waldstein, S. M. & Bogunovic, H. Artificial intelligence in retina. Prog. Retin. Eye Res. 67, 1–29, https://doi.org/10.1016/j.preteyeres.2018.07.004 (2018).

19. Dubis, A. M. et al. Democratizing Deep Learning Research Through Large Publicly Available Datasets and Tools. Investig. Ophthalmol. & Vis. Sci. 62, 1809–1809 (2021).

20. Arikan, M. et al. Uncertainty-based Deep Active Learning for Retinal Layer Segmentation. Investig. Ophthalmol. & Vis. Sci. 62, 2554–2554 (2021).

21. Li, J. et al. Multi-scale gcn-assisted two-stage network for joint segmentation of retinal layers and discs in peripapillary oct images. Biomed. Opt. Express 12, 2204–2220, 10.1364/BOE.417212 (2021).

22. Hassan, T., Akram, M., Masood, M. & Yasin, U. Deep structure tensor graph search framework for automated extraction and characterization of retinal layers and fluid pathology in retinal sd-oct scans. Comput. Biol. Medicine 105, 10.1016/j.compbiomed.2018.12.015 (2018).

23. Hassan, T., Akram, M. U., Werghi, N. & Nazir, M. N. Rag-fw: A hybrid convolutional framework for the automated extraction of retinal lesions and lesion-influenced grading of human retinal pathology. IEEE J. Biomed. Heal. Informatics 25, 108–120, 10.1109/JBHI.2020.2982914 (2021).

24. He, Y. et al. Retinal layer parcellation of optical coherence tomography images: Data resource for multiple sclerosis and healthy controls. Data Brief 22, 601–604, https://doi.org/10.1016/j.dib.2018.12.073 (2019).

25. Antony, B. J. et al. Automated segmentation of mouse oct volumes (asimov): Validation clinical study of a light damage model. PLOS ONE 12, 1–17, 10.1371/journal.pone.0181059 (2017).

26. Gholami, P., Kuppuswamy Parthasarathy, M., Roy, P. & Lakshminarayanan, V. OCTID citation. 10.5683/SP2/W43PFI (2018).

27. Farsiu, S. et al. Quantitative classification of eyes with and without intermediate age-related macular degeneration using optical coherence tomography. Ophthalmology 121, 162–172, https://doi.org/10.1016/j.ophtha.2013.07.013 (2014).

28. Tian, J. et al. Real-time automatic segmentation of optical coherence tomography volume data of the macular region. PLOS ONE 10, 1–20, 10.1371/journal.pone.0133908 (2015).

29. Chiu, S. J. et al. Kernel regression based segmentation of optical coherence tomography images with diabetic macular edema. Biomed. Opt. Express 6, 1172–1194, 10.1364/BOE.6.001172 (2015).

30. Chiu, S. J. et al. Validated Automatic Segmentation of AMD Pathology Including Drusen and Geographic Atrophy in SD-OCT Images. Investig. Ophthalmol. Vis. Sci. 53, 53–61, 10.1167/iovs.11-7640 (2012). https://arvojournals.org/arvo/content_public/journal/iovs/932974/z7g00112000053.pdf.

31. Melinšcak, M., Radmilovic, M., Vatavuk, Z. & Loncaric, S. Annotated retinal optical coherence tomography images (aroi) database for joint retinal layer and fluid segmentation. Automatika 62, 375–385, 10.1080/00051144.2021.1973298 (2021). https://doi.org/10.1080/00051144.2021.1973298.

32. Morales, S. et al. Retinal layer segmentation in rodent oct images: Local intensity profiles fully convolutional neural networks. Comput. Methods Programs Biomed. 198, 105788, https://doi.org/10.1016/j.cmpb.2020.105788 (2021).

33. Rasti, R., Rabbani, H., Mehridehnavi, A. & Hajizadeh, F. Macular oct classification using a multi-scale convolutional neural network ensemble. IEEE Transactions on Med. Imaging 37, 1024–1034, 10.1109/TMI.2017.2780115 (2018).

34. Hitachi-Automotive-And-Industry-Lab. Semantic segmentation editor. https://github.com/Hitachi-Automotive-And-Industry-Lab/semantic-segmentation-editor (2022).

35. Willoughby, J. et al. Object detection on medical images with the aid of contrastive gated attention. Investig. Ophthalmol. & Vis. Sci. 63, 2998 – F0268–2998 – F0268 (2022).

36. Kermany, D. S., Zhang, K. & Goldbaum, M. H. Large dataset of labeled optical coherence tomography (oct) and chest x-ray images (2018).

37. Tan, M. & Le, Q. V. Efficientnet: Rethinking model scaling for convolutional neural networks (2020). 1905.11946.

38. Yakubovskiy, P. Segmentation models. https://github.com/qubvel/segmentation_models (2019).

